# Dynamics of a Form-Fitting Protein in a Nanopore: Avidin in ClyA

**DOI:** 10.1101/221457

**Authors:** Bo Lu, Chris Stokes, Monifa Fahie, Min Chen, Jene A. Golovchenko, Lene Vestergaard Hau

## Abstract

We probe the molecular dynamics of a protein, avidin, as it is captured and trapped in a nanopore, ClyA, with time-resolved single-molecule electrical conductance measurements, and we present a method for visualizing this process from the data. The case of avidin in ClyA has rich time-dependent conductance spectra of discrete levels that correlate with different configurations of the protein in the pore. One is very long-lasting, stable and noise-free, and portends the use of this system as a platform for more general studies of proteins and other molecules, where avidin acts as a shuttle that ferries analytes into the pore for probing. We demonstrate this by the sensitive detection of a biotin molecule attached to avidin captured by the pore. We also present an approach to determining the nanopore size based on a 3D printed model of the pore.

## INTRODUCTION

It is possible to observe individual charged biological molecules as they translocate through a voltage biased nanopore in a lipid membrane by monitoring the ionic conductance of the pore during the molecular motion through the pore. Such measurements have provided new insights into the biophysics of these molecules and their interactions with the pores^1^. They have also resulted in practical, portable instruments for sequencing DNA^2^. Probing proteins by this translocation method has provided interesting insights but proved challenging due to the more complex geometrical and charge structures of proteins, however progress is being made^3-8^.

We have been particularly inspired by recent nanopore studies^9,10^ where protein molecules are transiently trapped and electrically monitored in the large *ClyA* nanopore. In the following we show that when a very close match between the protein size and the pore lumen size is obtained, remarkably detailed, time-dependent electrical-conductance spectra are observed that reveal discrete levels including low-noise and long-lasting trapped states that can be cleared deterministically with a control voltage. We show that these spectra, appropriately analyzed, provide valuable insight into protein and pore structure and the dynamical landscape of their mutual interactions. We conclude that a judiciously chosen protein-pore pair can serve as a platform for the study of a wide range of other molecules, including proteins and their interaction with substrates in solution.

The size-matched pair we have discovered is avidin and *ClyA*_12_, a dodecamer nanopore. Avidin is a positively charged protein with a high isolelectric point of roughly 10^11,12^, and its size is well understood from X-ray studies^13, 14^ The *ClyA* pore on the other hand exists in several oligomeric forms^15^, each having a different pore size and open-pore conductance. We identify the most common pore that we observe as the dodecamer, *ClyA*_12_, with charge separation experiments on the relevant ClyA nanopore and conductance measurements on a 3D printed macroscopic analog. Finally we demonstrate the utility of the new nanopore platform with the sensitive detection of biotin attached to avidin proteins that are captured and ejected from a single pore.

## RESULTS

Figure 1a shows molecular models, with dimensions, of the avidin and dodecamer *ClyA*_12_ nanopore. The close match between the avidin outer dimensions and the pore lumen’s inner dimension makes this pair attractive for our purposes. These pictures and dimensions have been determined by X-ray diffraction^13,14,16^ and can be found in the protein database^17^. (More detailed descriptions of ClyA and avidin are in Online Methods.)

**Figure 1.**
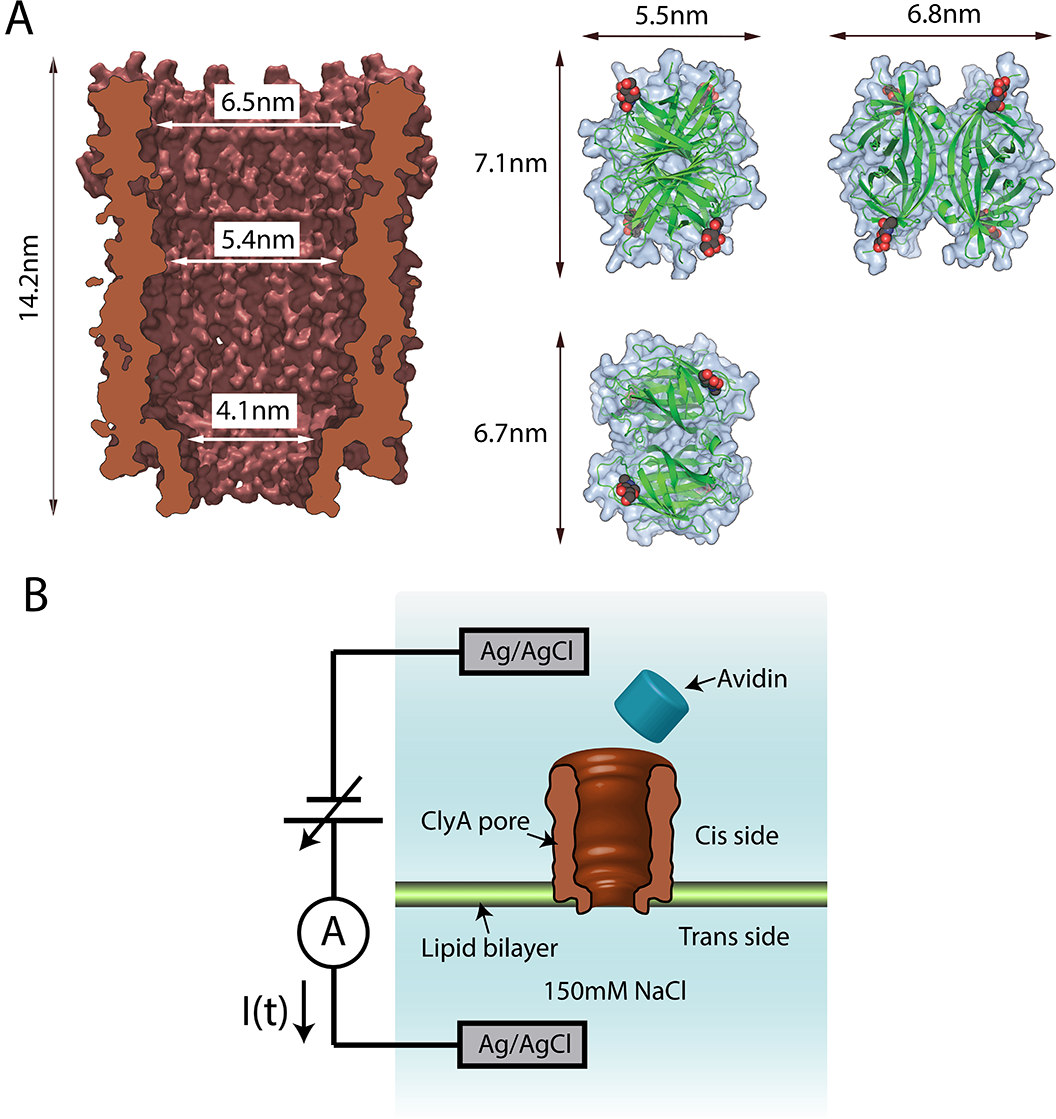
Avidin-ClyA Setup. **(a)** Protein structures of the ClyA dodecamer pore and of avidin. The figures are based on the x-ray crystallography structures 2WCD and 1AVE, respectively, from the Protein Database. The ClyA pore is shown in a cross-sectional view along the center plane containing the twelve-fold rotational symmetry axis. Avidin is viewed from three different directions. Two of its four beta barrels are visible in the figures (green) as are two of the four N-Acetylglucosamines (oxygens are red). **(b)** Experimental setup. A lipid bilayer membrane is suspended across a 40 micrometer teflon frame in an electrolytic buffer solution (150mM NaCl, 15 mM TRIS pH 7.5). A silver/silver-chloride electrode is placed in each reservoir and a voltage of a few tens of mV, typically, is applied across the membrane. The membrane electrically isolates the two reservoirs, and initially no current flows between the electrodes. Preformed ClyA pores are added to the cis side, and when a single pore inserts in the membrane – observed as a step change in current to a stable open-pore current – the cis side is immediately flushed with buffer electrolyte solution to prevent additional ClyA pores from inserting. Once a single pore is in place in the membrane, 10 pmol of avidin is added to the cis chamber. With a negative bias voltage applied, the current is transiently blocked when individual, positively charged avidin molecules are trapped by and subsequently escape or, by voltage reversal, are ejected from the pore.

### Experimental Setup

Figure 1b illustrates the experimental setup. Two chambers of electrolyte solution (150mM NaCl, 15 mM TRIS at PH 7.5) are separated and electrically isolated from each other by a lipid bilayer of 1,2-diphytanoyl-*sn*-glycero-3-phosphocholine (Avanti Polar Lipids, Inc.) stretched across a 40 micron diameter teflon frame. Ag/AgCl electrodes are inserted in each chamber and a voltage bias is applied across the membrane. For later trapping of positively charged avidin in the pore, the trans chamber is kept at negative voltage. ClyA nanopores are first added to the cis chamber. When a single pore inserts in the membrane, the ionic current induced by the voltage bias abruptly increases after which the remaining nanopores in solution are removed (flushed) from the cis chamber.

The current observed for individual voltage-biased nanopores is measured, and for the work reported here, nanopores with a conductance of 1.66 nS (± 1%) at 30 mV bias were selected. Pores with this conductance value are the most commonly observed in the membrane and they are the most stable over time. (For experimental details, see Methods.)

### Avidin in a 1.66 nS pore

After 10 pmol of avidin is added to the 250 microliter cis chamber with a 1.66 nS pore in place, the current through the – 35 mV voltage biased pore is observed to transiently drop from the open pore value (58 pA) as individual avidin protein molecules are captured by, and escape from, the pore (Fig. 2a-g). These current blockage events are separated into two categories. The first we call Transient Captures. It consists of transient current blockage events that last from 200 microseconds up to 1 sec before the pore current returns to the open-pore value after a captured protein has spontaneously escaped from the pore. The second category we call Permanent Captures. These events last for times exceeding 1 second, and the captured protein is almost never observed to spontaneously leave the pore. So for these events we automatically reverse voltage bias after one second of current blockage. This ejects the positively charged protein from the pore. The voltage bias is then returned to a negative value and the open pore current is again observed, followed by new current blockage capture events. Thus our Permanent Events all last for 1 second at which time the trapped avidin is forcibly ejected from the pore.

**Figure 2.**
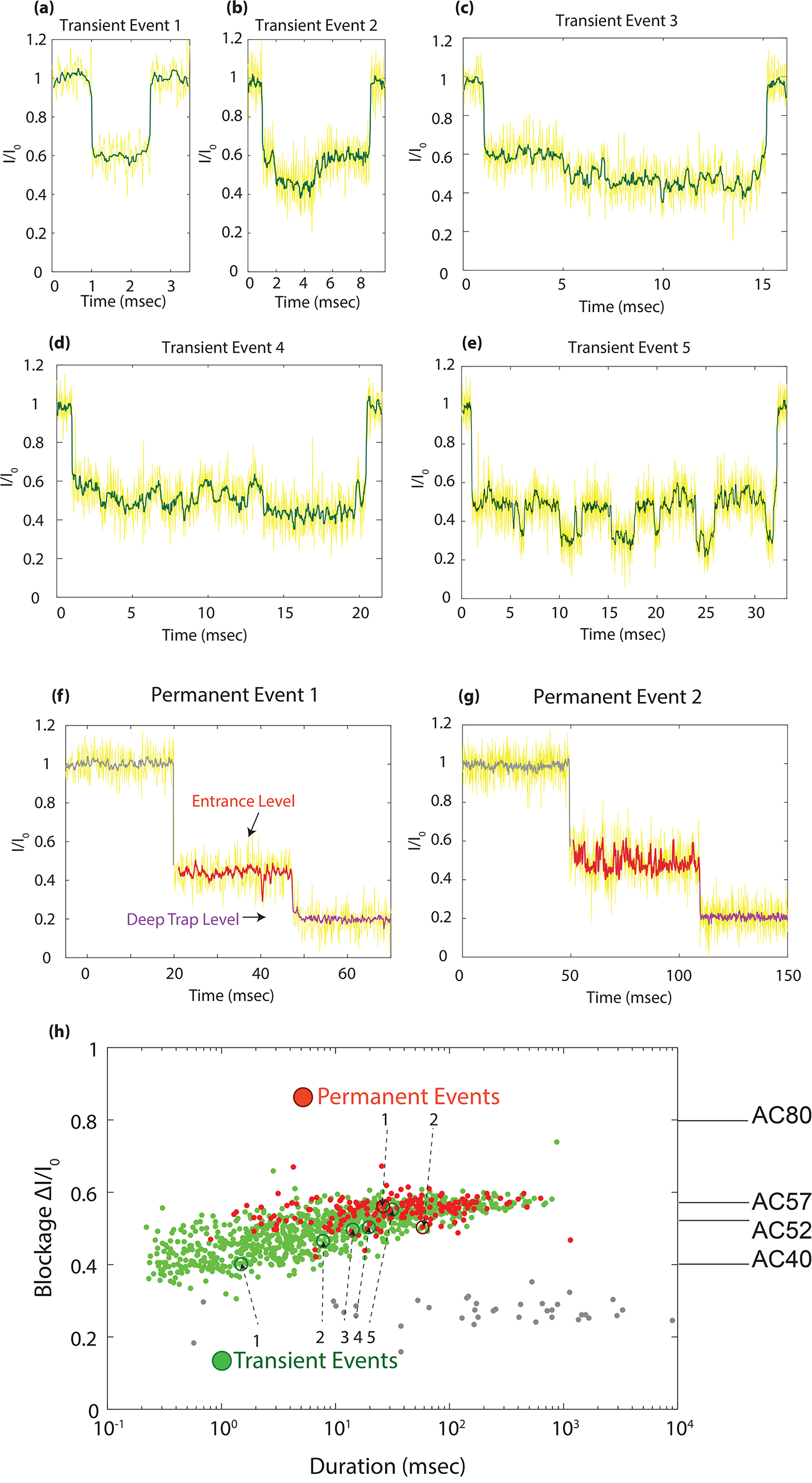
Capture events for individual avidin molecules. **(a-g)** Time traces for capture events. The first five **(a-e)** are transient events of increasing duration. Some have a rich time structure, with discrete blockage levels and transitions between them. The last two time traces **(f,g)** are for permanent events. Here, an immediate current drop to an entrance blockage level is followed by a transition to a deep, permanent capture level with very low current fluctuations. For all permanent events, we apply a +100 mV voltage pulse to eject the avidin from the pore after 1 second of trapping. **(h)**. Scatter plot of event blockage versus duration. Each transient event, shown as a green dot in the figure, is characterized by its duration and average blockage. Each permanent event, represented by a red dot, is characterized by the time spent at the entrance level, and the corresponding average blockage, before transition into the permanent, deep blockage level, AC80. The seven events in **(a-g)** are indicated in the figure. We also mark the blockage levels AC40, AC52, and AC57 discussed in the text and visible in one or more of the time traces in **(a-g)**.

Each single protein capture, whether transient or permanent, is observed to have its own unique time-dependent current trace during the event. Figures 2a-g show a collection of single molecule capture events in the 1.66 nS pore. We plot the observed current *I*(*t*) during each event, divided by the open pore current *I*_0_. The events in Figs. 2a-e show the time-dependent blockage for transient events of increasing duration. Most transient events are simple, having only a single blockade level as in the first event in Fig. 2a. The longer transient events, Fig. 2b-e, have been chosen to show that sometimes transitions between levels are observed during a single event. In a first-pass analysis, each Transient Capture is characterized by two numbers: the duration and the average blockage during the event. The “blockage” is defined to be (*I*_0_ – *I*(*t*))/*I*_0_ averaged over each event’s duration. Figure 2h is a “scatter plot” of these parameters for each Transient Capture event (green points) reflecting this characterization. Each green point belongs to one protein capture. Scatter plots for other bias voltages are shown in SI Fig. 1.

As the Transient Capture event duration increases from a few hundred microseconds to a few hundred milliseconds, the average blockage slowly increases. In the next figure we will show that the reason for this is the existence of multiple discrete blockage levels that are averaged over in each event in the scatter plot. A few examples of separate and clearly distinguishable blockage levels and transitions between them are already seen in Figs. 2a-e. These events are also numbered in the scatter plot. Shortly, we will present a quantitative and global view of these complex dynamical signals within the event distribution.

Figures 2f-g show the current traces for two Permanent Captures. Each of these events has an interesting time structure. As with most Permanent Capture events they start with a preliminary, fluctuating blockage that is followed by a deep and quiet permanent blockage level. Permanent Captures are characterized by the time duration and average blockage during the preliminary part of the event. With these parameters, each Permanent Capture event is added to the scatter plot of Fig.2h (red). The average blockage is clearly independent of duration and close to that of the longest transient events. This immediately suggests that, prior to capture to a deep permanent blockage level, the protein often passes through an intermediate state of variable duration. These permanent blockages are of special importance for future protein studies. This will be discussed later. They often persist even after the voltage bias has been removed. They can always be cleared by reversing the bias voltage.

### Protein Dynamics Landscape Plots

The existence of discrete blockage levels and their time-dependent populations is shown much more dramatically and quantitatively in the protein dynamics landscape (PDL) plots introduced in Fig. 3. To generate a particular histogram in Fig. 3a, only transient events that last for times less than a specified ***τ***_*trans*_ are included. Then the amount of time spent (in units of 160 microseconds) at each blockage level for all these events is determined and a histogram of the time spent at each level is generated. We refer to this histogram as a blockage spectrum. A collection of such blockage spectra, for a set of increasing ***τ***_*trans*_ values, is a Protein Dynamics Landscape.

**Figure 3.**
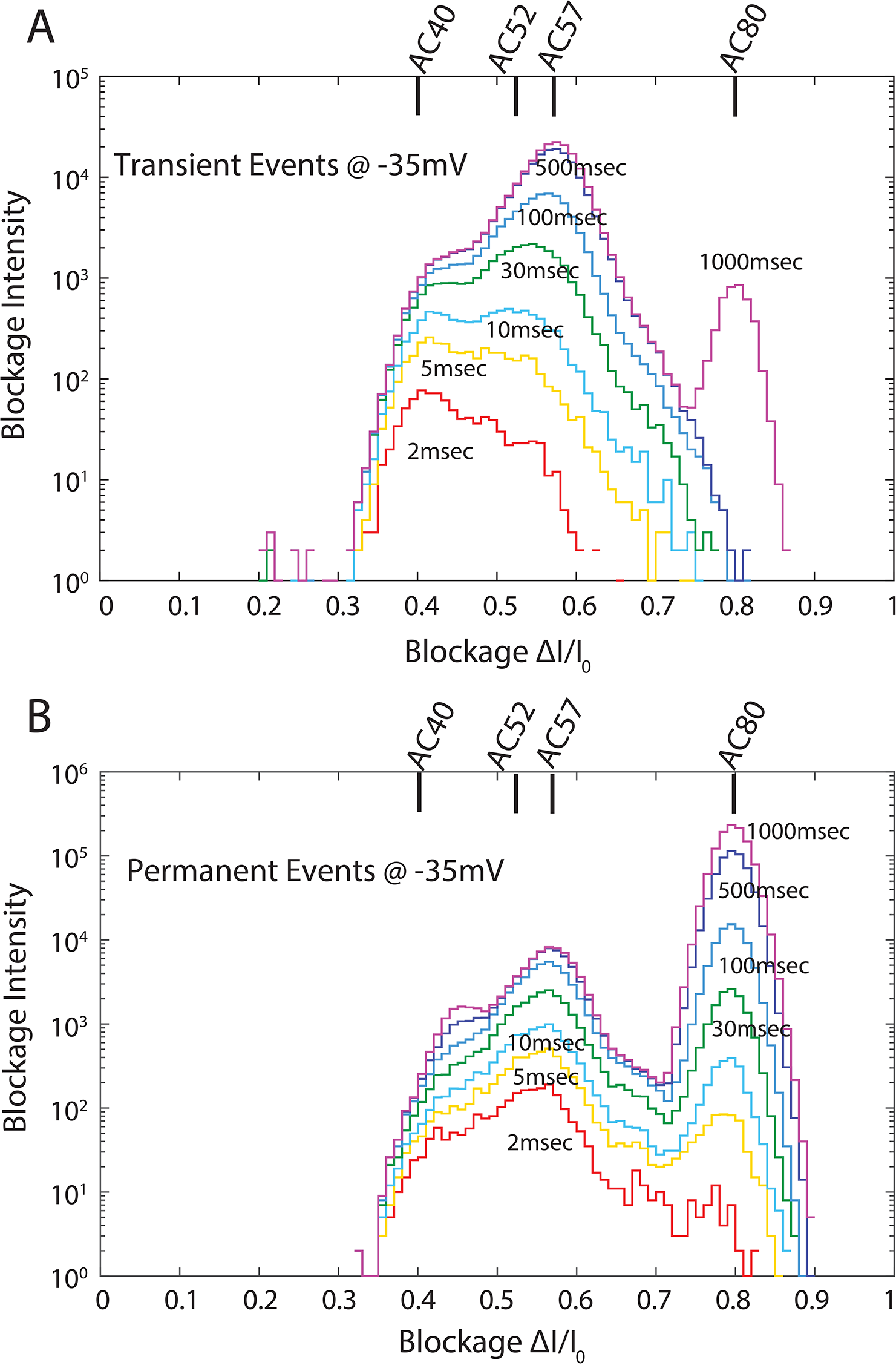
Protein Dynamics Landscape (PDL) Plots. **(a)** Transient events. Individual current blockage spectra are first obtained by grouping together all transient events that last for less than a given time, ***τ***_*trans*_, and then generating a histogram of the number of times each blockage level appears when averaged over a 160 microsecond window bin. Then the plot is generated from a whole set of such histograms of level populations, obtained by varying the value ***τ***_*trans*_ from 2 ms to 1000 ms, as indicated in the figure. **(b)** Permanent Events. Here all (205) permanent events are grouped, and each histogram refers to the number of times a blockage level appears up until a time ***τ***_*perm*_ after the capture events start. The plot is generated from a set of histograms with ***τ***_*perm*_ varying from 2ms to 1000 ms. For both plots, the voltage is fixed at −35 mV. The discrete blockage levels shown at the top of each plot are deduced directly from these plots by non-linear least-squares fitting with sums of Gaussians, as described in the Supplementary Information.

The PDL here is obtained for a fixed bias voltage of −35 mV. The plot reveals several peaks, corresponding to discrete blockage levels for avidin in ClyA. Note that deeper blockage levels last longer than shallower levels.

A similar analysis can be performed on permanent capture events and the results are shown in Fig. 3b for the same bias voltage of −35 mV. Here, all permanent capture events are considered together and each histogram refers to the accumulated time (in units of 160 microseconds) a blockage value appears up until ***τ***_*perm*_ after the capture events start (which is when avidin first enters the pore). The deepest blockage level is maximally populated for the longest times.

A complete set of PDL plots is obtained from data like those of Figs. 3a-b at different bias voltages and are shown in Supplementary Information (SI) as Fig. SI 2.

In the PDL plot of Fig. 3a, we see a single peak centered at 40% blockage for transient events lasting 2 ms or less. We call this level (Avidin Capture) AC40. As longer events are included (i.e., as ***τ***_*trans*_ is increased), a second peak emerges. With careful fitting, we determine this level to be centered at 52% blockage and denote the level AC52. At 10 ms the two peaks have equal intensity, and at 30 ms, the AC52 peak dominates. For longer times, a separate (AC57) peak at 57% blockage dominates, and finally we observe a few very deep captures for transient events that last longer than 1/2 second. Here the blockage level at 80% dominates (AC80). Gaussian fits to the spectra clearly reveal a fifth peak at 45% blockage (AC45), and the fits are shown in Fig. SI 3.

It is interesting to compare this dynamical behavior to that revealed by the PDL plot for permanent events at −35 mV (Fig. 3b). During the first 2 ms of the permanent events, the spectrum is dominated by one peak: AC57. The AC40 peak is barely visible, and AC52 is missing. The spectra are generally dominated by AC57 during the early times of the permanent events, but later (>=100 ms) the AC80 peak dominates.

We conclude that for the permanently trapped events, avidin is first captured into state AC57, where it stays for an average of 60 ms (at −35 mV). This is followed by capture into a deeply and permanently trapped state, AC80. This picture is supported by the time traces for the Permanent Capture events in Figs. 2f-g.

We have found that the level populations in the PDL’s are strongly voltage-bias dependent (Fig. SI 2). This effect is already dramatically seen by simply plotting the ratio of the number of Permanent to Transient events as a function of bias voltage. This is shown in Fig. 4. Increasing the bias voltage from 30 to 50 mV results in dramatic and exponential increase in this ratio. At high bias voltage, the energy landscape of the protein in the pore is clearly biased towards permanently capturing the avidin to the deepest level in the pore from which escape eventually becomes impossible.

**Figure 4.**
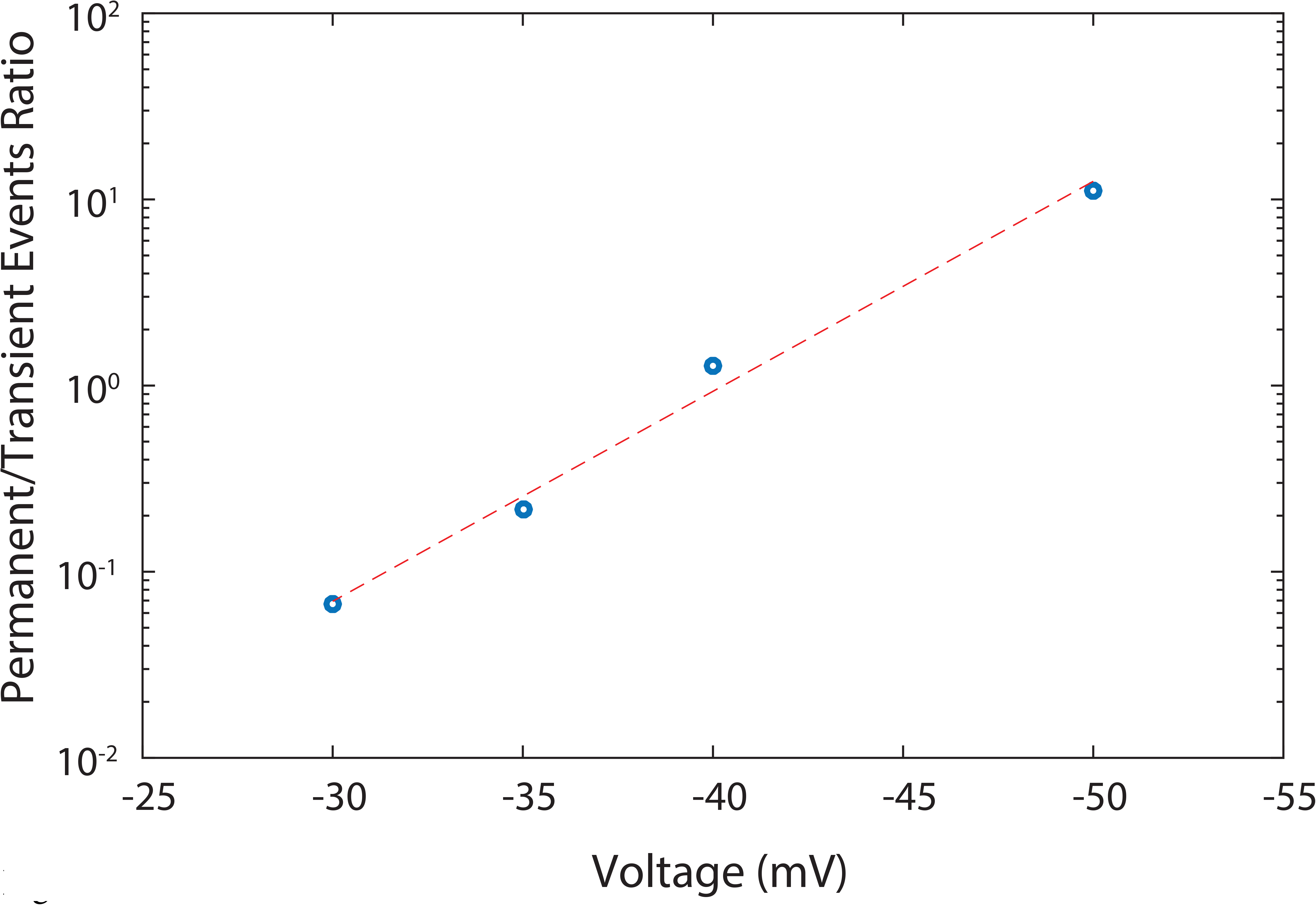
Ratio of the number of permanent to escape events as a function of voltage.

### ClyA_12_ Pore Conductance Modeling

The avidin-ClyA experiments described above for a 1.66 nS pore can ultimately only be understood on a molecular level if the number of protomers in the ClyA oligomer is known. To address this issue we have devised a series of experiments to determine which oligomer is expected to have the observed conductance of 1.66 nS. The problem is complicated by the fact that there are two phenomena that determine the conductance. One is simply the geometry of the pore. This geometrical conductance is however modified by the presence of charged amino acids on the walls of the nanopore. In fact, in ClyA the limiting aperture is known to have a highly negative charge which can block chloride ion conductance through the pore^18^. We now show that geometric and charge selectivity effects can be determined in different experiments and assembled to predict the total conductance of a given nanopore.

To determine the geometrical contribution to the pore conductance, we constructed a 3D printed model of a *ClyA*_12_ pore scaled up by a factor of 10^7^/3 to atomic coordinates for this pore obtained from the protein database. This macroscopic pore was glued into a thin insulating plastic sheet to simulate the membrane. The sheet was used to separate two reservoirs of 150 mM NaCl in water, and in each reservoir hemispherical electrodes of 11.4 cm radius surrounded the pore. Further details are in Methods and a schematic of the experiment is in Fig. SI 4.

Scaling the measured conductance value by 3/10^7^ to obtain the geometrical conductance for the nanopore gives 2.87 nS. (The inverse of the measured geometric pore conductance is a sum of the inverse conductance for the pore alone, and the access resistance^19^. The correction of access resistance to infinite electrode radius increases the total resistance by 1.57 per cent, and this correction has been included.)

This measured and scaled conductance value will be significantly reduced by exclusion of part of the chloride ions’ contribution to the pore conductance. To evaluate the reduction factor we captured a 1.66 nS pore in a membrane with a 150 mM buffered salt solution on both sides. The cis side of the membrane was then refilled with a 20 mM buffered salt solution. This resulted in a measurable open circuit potential of 26 mV across the pore due to the unequal transfer of sodium versus chloride ions across the pore in the drift-diffusion process.

The GHK equation from ion-channel biophysics^20^ is designed to handle this situation, and with the measured open circuit potential across the pore, we can use the equation to calculate the permeability factor reflecting the extent to which the chloride ions are blocked, relative to the sodium ions, from passing through the pore. We obtain a permeability factor of 0.22, which is smaller than the 0.33 obtained at higher salt concentrations^18^. We would indeed expect a larger chloride ion blockage effect in our experiments due to a larger Debye screening length at lower ionic strength. The obtained permeability factor reduces pore conductance from the geometrical pore conductance value, and we predict a *ClyA*_12_ pore conductance under the conditions of our experiment to be 1.75 nS. This is in excellent agreement with the measured value of 1.66 nS. Simple scaling arguments show that the conductance prediction for the other possible oligomer candidate, the slightly larger 13’mer, would be 1.97nS, which rules it out. Note however that we have observed a small number of pores with 1.9 nS but they are somewhat unstable when inserted in the membrane.

### 3D Printed Protein in a 3D Printed Pore

We prepared a 3D printed avidin protein with the same scaling as used for the *ClyA*_12_ 3D printed pore. Regardless of orientation, the protein was slightly too large to completely enter the pore lumen, mostly being blocked from entry by a few residues on the outside surface of the avidin. Placing the molecule on the rim of the pore in the macroscopic conductance experiment (Fig. 5a) only reduced the conductance by 10%. With avidin deeply inserted in the lumen of the pore (Fig. 5b), only a 20% conductance reduction was observed. We postulate that a full understanding of the much larger conductance decreases for the 1.66 nS nanopore experiment may also be sought in a geometrical and/or charge exclusion explanation. Geometrically, elasticity of the protein and/or the pore may allow the protein to enter into the pore more deeply and with more intimate contact under the force provided by the electric field from the applied voltage bias. In addition, electrostatic charges in the pore lumen may interact strongly with electrostatic charges on avidin at short distance when avidin is near the bottom of the pore. And ion charge-selectivity effects for conductance may become more intense when the charged protein is in the pore, also leading to deeper current blockades. All effects are of great interest and accessible to future research.

**Figure 5.**
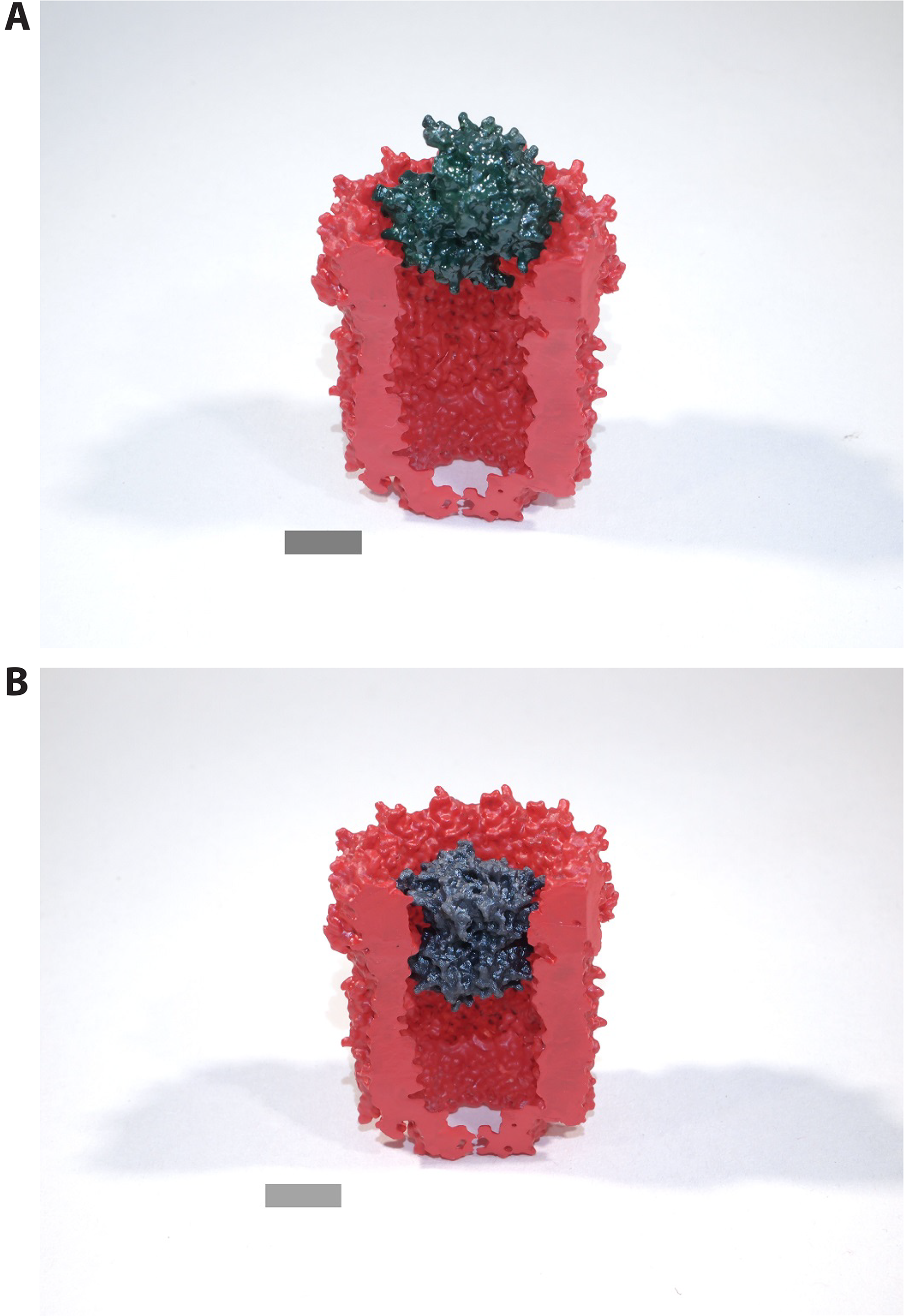
3D printed macroscopic models. **(a)** Rigid avidin model atop rigid *ClyA*_12_ pore oriented to obtain maximum current blockage of 10% in macroscopic conductivity experiment. **(b)** Elastic avidin model maximally pressed into rigid *ClyA*_12_ pore model achieving 20% blockage in macroscopic conductivity experiment. Scale bar corresponds to 3 nanometers for actual protein and nanopore.

### Biotin-Avidin Complex in a 1.66 nS pore

Finally we demonstrate the sensitivity of the avidin-ClyA platform for detecting other molecules attached to avidin, Here 100 pmoles of biotin was added to the cis chamber already containing 10 pmoles of avidin at −35 mV voltage bias. After a few minutes we observed a dramatic change in the trapping dynamics. The ratio of permanent to transient event rates was reduced by a factor of 28, from 1/6.4 (avidin) to 1/182 (biotin-avidin) with the latter closely resembling the ratio for avidin alone at -30mV bias voltage (1/159). From this we conclude that the net positive charge of the biotin-avidin complex is lower by a factor of 30/35 compared to avidin alone. Calculations^21^ based on the PDB crystallographic structures for deglycosylated avidin (1AVE) and for the complex of deglycosylated avidin with four biotin ligands (2AVI) predict a charge of the biotin-avidin complex that is lower by 2 positive charge units than that of avidin alone. (Note, in both these cases avidin is deglycosylated except for the core glucosamine (GlcNAc).) Combining this with our observations, we obtain a charge of +14e for fully glycosylated avidin (used in our experiments).

Interestingly, we also observe that for the biotin-avidin complex, the current blockage for the deeply trapped level is decreased by 4% relative to that for avidin alone. This is shown in Fig. 6. Since the deep blockage level for avidin is independent of voltage in the -30 to −35 mV range, we posit that the difference in deep blockage levels is due to a slightly bigger size or lower elasticity of the biotin-avidin complex^13,22^. An alternative explanation is a weaker electrostatic attraction of biotin-avidin to the negative charges at the limiting aperture of ClyA.

**Figure 6.**
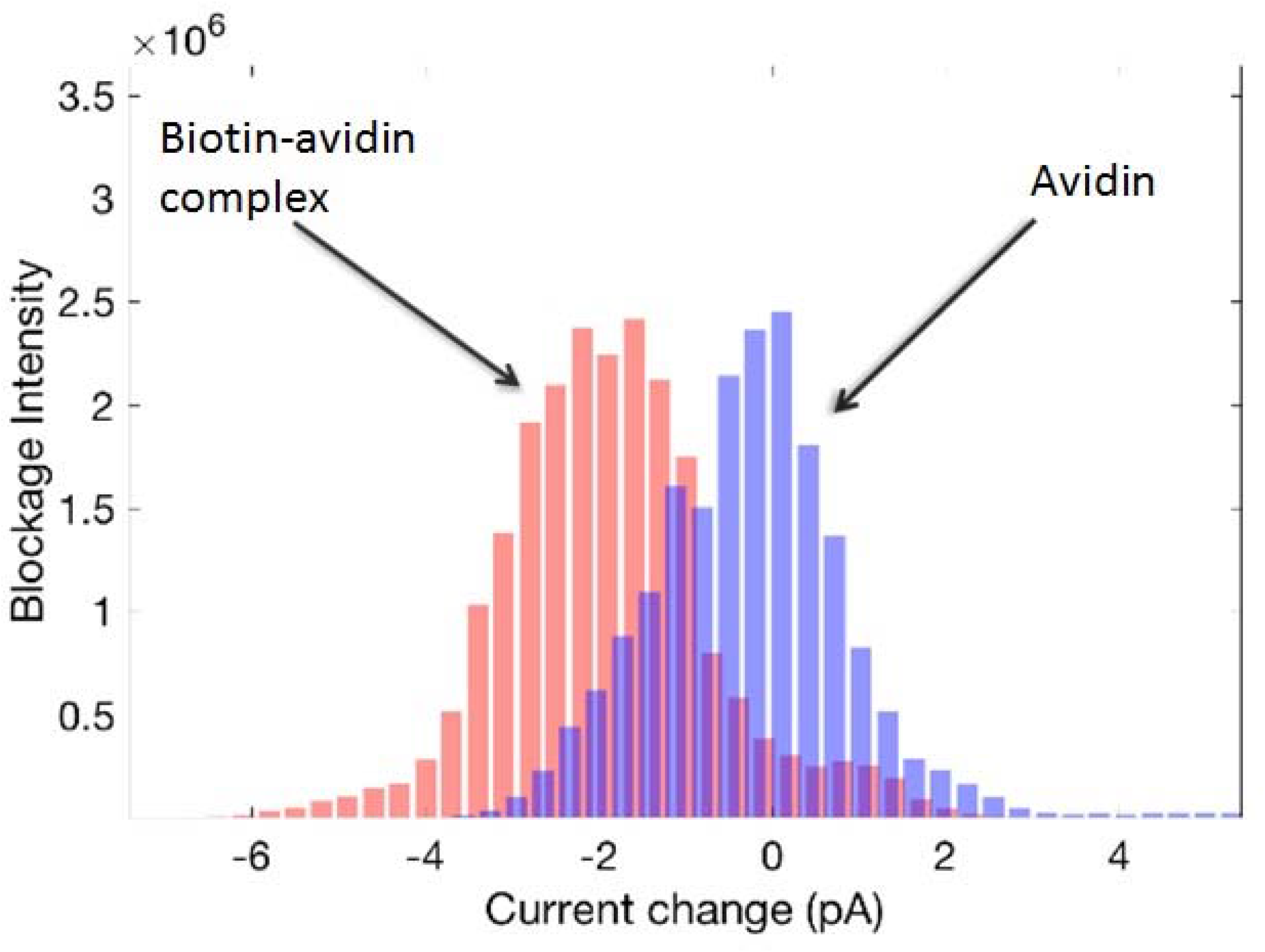
Biotin-avidin complex. Current blockage spectra for the deeply trapped level for both the biotin-avidin complex and for avidin alone in a *ClyA*_12_ nanopore biased at −35mV. The unit for the vertical axis is 4 microseconds. A 100-point median filter has been applied to the raw data sampled every 4 microseconds.

## DISCUSSION

Our experiments and analysis show that the avidin-ClyA nanopore system can be electronically probed and quantified to a remarkable degree. Discrete conductance levels, transitions between them, and the fluctuations within them all reflect the interaction between, and time evolution of, avidin and the voltage biased pore in an electrolyte environment.

One may expect that the qualitative discussions we have presented will be made more precise when the powerful computational tools of molecular modeling and simulation of ionic conduction in confined spaces are applied to this system.

An important reason for focussing on the avidin-ClyA system is its potential as a new platform for the study of protein structure, dynamics, and enzymatic activity at the single molecule level. It is already standard practice to attach other biomolecules to avidin-like proteins via biotin linkers^23,24^, and the avidin will allow nanopore capture, for a controlled capture time, of a biotin-linked protein, facilitating study of the attached protein. A first step in this direction has already been taken by our detection of the small molecule biotin attached to a captured avidin protein.

## METHODS

Methods and any associated references are available in the online version of the paper.

## ACKNOWLEDGMENTS

We thank Eric Brandin for biochemical preparations and evaluations and Stephen Fleming for assistance and advice.

## AUTHOR CONTRIBUTIONS

LB contributed experimental work and computing assistance at Harvard, CS prepared the 3D protein models and performed experiments at Harvard, MF prepared ClyA at Amherst, MC directed work on expression and purification of ClyA at Amherst, JAG and LVH conceived the idea for the experiment and method of analysis, and directed research at Harvard, and in addition LVH conducted experiments at Harvard.

## ONLINE METHODS

### The Protein – Nanopore System

Cytolysin A (*ClyA*) is a pore forming cytolytic toxin expressed in several pathogenic strains of *Escherichia coli* and *Salmonella enterica*. Upon reaching a target-cell membrane, the soluble 34 kDa ClyA monomers undergo significant conformational changes, involving half of the monomers’ amino acid residues, and subsequently assemble into membrane-bound oligomers^16,25^. The resulting membrane-spanning *ClyA* pores form nanometer-sized circular holes in the membrane whose function as a cell barrier is lost, and this ultimately leads to cell death. It is clear from multiple observed pore conductances and gel studies that different oligomers of ClyA exist and may be of use for other protein nanopore studies^15^.

Commercially available avidin (Pierce Avidin, ThermoFisher Scientific) is purified from hen egg white, and the molecule is a tetramer that consists of four beta barrels. Each barrel is open at one end, which allows one end of its ligand biotin to enter and bind to residues inside the beta barrel^13,14^. (In Fig. 1a, two of the beta barrels (green) are clearly visible.) The binding of biotin to avidin represents the strongest non-covalent ligand bond found in nature (with a dissociation constant of 10^−15^M), and the protein-ligand pair has been used extensively in molecular biology, bio-technology, and medical applications^23^. In these experiments, we first use the apo-avidin (with no biotin bound). Avidin is a glyco-protein with one asparagine glycosylation binding site per monomer (Asn17). A single core N-Acetylglucosamine (GlcNAc) is visible at two sites in the image which is based on the crystal structure 1AVE in the protein database^17^. This structure has a charge of roughly 7 positive charges at the pH of 7.5 used in the experiments in this paper^21^.

Avidin used in our experiments (Pierce Avidin, ThermoFisher Scientific) has a polysaccharide attached to each of the four Asn17 locations. In addition to the core GlcNAc, the polysaccharides have 4–5 mannose and 2 GlcNAc^12^.

### Picoamp-Current Measurements

For all current measurements presented here, we used a Molecular Devices Axopatch 200B patch clamp amplifier. The output signal was processed by a 10 kHz, 4 pole Bessel filter to minimize high frequency current noise. After filtering, the signal was sampled and digitized every 4 microseconds and recorded in computer memory for further processing and analysis.

### Macroscopic 3D Models and Conductance Modeling

The 3D printer used to prepare the scaled-up protein and nanopore was a FormLabs Form 2 model, with a resolution of ~50 microns. The dimensions of the molecules were scaled up by a factor of 10^7^/3 from the PDB database. The plastic used for the rigid models was FormLabs Clear (Part # FLPGCL02, FLPGCL03). The plastic used for the flexible avidin model was FormLabs Flexible (Part # FLFLGR02) with a Shore Hardness of 80A.

Conductance measurements on the macroscopic pore system were made with a range of 60 Hz AC voltages from 10-50 volts rms. Conductance was calculated from the linear slope of the I-V plot. No phase shifts between observed currents and applied voltages were observed. Temperature of the electrolyte was monitored and measurements were restricted to a few seconds in duration to minimize electrolyte Joule heating.

### ClyA Monomer Expression and Purification

All reagents were purchased from Fisher Scientific and/or Boston Bioproducts unless otherwise stated. Phenylmethane sulfonyl fluoride (PMSF) and magnesium chloride were purchased from Sigma.

C-terminal His6 tagged ClyAwt protein was expressed in BL21 (DE3) cells. Specifically, pT7-ClyAwt-CHis6 plasmid was transformed in BL21 (DE3) chemically competent cells and grown on LB-Amp Agar plates. One colony was inoculated in starter LB media containing 100 μg/ml ampicillin antibiotic and grown at 37°C with shaking at 200 rpm. The starter culture was used to inoculate 250 ml LB media containing 100 μg/ml ampicillin. The culture was grown at 37°C until the OD_600_ was between 0.5 and 0.65. The culture was then cooled on ice and induced by adding IPTG to a final concentration of 0.5 mM and then incubated for 16 hrs at 15°C with shaking. After 16 hrs, the culture was harvested at 3100 × g and the pellet resuspended in 15 ml of 50 mM Tris-HCl pH 8.0, 1 mM EDTA buffer and frozen in -20°C until ready to use.

The frozen pellet was subsequently thawed at room temperature and a final concentration of 0.5 mM PMSF was added. The mixture was sonicated on ice to lyse the cells. MgCl_2_ was added to the lysate at a final concentration of 10 mM and the mixture was then centrifuged for 20 mins at 20,000 x g. The supernatant was filtered through a 0.22 μm filter membrane and loaded onto a gravity NiNTA column equilibrated with buffer A (150 mM NaCl, 50 mM Tris-HCl pH 8). The column was subsequently washed with buffer A to remove unbound proteins. Buffer A1 (150 mM NaCl, 50 mM Tris-HCl, 50 mM imidazole) was used to wash the weakly bound proteins and then the ClyA protein was eluted and collected in buffer A2 (150 mM NaCl, 50 mM Tris-HCl, 150 mM imidazole).

The eluted ClyA proteins were dialyzed using a 6-8 kDa cutoff membrane with constant stirring at 4°C for two cycles in dialysis buffer (150 mM NaCl, 50 mM Tris-HCl, 5 mM EDTA). The proteins were then concentrated using a 10 kDa cutoff centricon to ~3 ml and loaded onto a gel filtration column equilibrated in 150 mM NaCl, 20 mM sodium phosphate pH 7.0 buffer to remove aggregated proteins. The ClyA monomer was collected and kept at 4°C for 2 weeks or in -80°C for long-term storage.

### Preparation and Purification of ClyA Oligomers (Nanopores)

Purified ClyA monomers were suspended at 0.6 mg/mL in a buffered solution containing 50 mM NaCl, 10 mM sodium phosphate pH 7.4 (with a buffer exchange column). Oligomeric ClyA was formed from monomers by the addition of n-Dodecyl beta-D-maltoside (DDM, Calbiochem/EMD Millipore; 10% w/v in water) to a final concentration of 1% and incubated 20 min at room temperature.

ClyA nanopore purification was carried out by blue native gel electrophoresis using a 4-16% polyacrylamide gradient gel (NativePAGE, Invitrogen/Novex Life Technologies). Typically, 10 ug of ClyA oligomers were combined with electrophoresis loading buffer and applied to a 1.0 mm X 5.0 mm sample well of the gel. Major bands of oligomeric ClyA were excised from the gel following electrophoresis, and nanopores were recovered from the gel slices by diffusion into an elution buffer containing 150 mM NaCl, 0.2% DDM, 50 mM Tris-Cl pH 8.0.

### Preparation of Avidin

Lyophilized purified avidin from hen egg white (Pierce/Thermo Scientific Product# 21121) was dissolved in deionized water to 2 mg/mL concentration. For subsequent storage at 4°C, an equal volume of 2X Phosphate Buffered Saline with 20% glycerol was added to the suspension to bring the avidin stock solution concentration to a nominal 1 mg/mL. Prior to use in ClyA nanopore experiments, an aliquot of the avidin stock solution was applied to a Bio-Spin 30 spin column (Bio-Rad Laboratories) equilibrated with 150 mM NaCl, 15 mM Tris-Cl pH 7.5 for buffer exchange.

### Preparation of d-Biotin

Biotin was prepared with 0.2 mg d-Biotin (Sigma-Aldrich/Millipore Sigma) per mL of 20 mM KCl, 50 mM Tris-Cl pH 7.6, and diluted in 150 mM NaCl, 15 mM Tris-Cl pH 7.5 to a final biotin concentration of 100 μM.

